# Chromosome-level Genome Assembly of the South African Lion (*Panthera leo melanochaita*)

**DOI:** 10.64898/2026.03.10.710750

**Authors:** Sibusiso Hadebe, Thendo Stanley Tshilate, Nompilo Hlongwane, Lucky Tendani Nesengani, Sinebongo Mdyogolo, Annelin Molotsi, Rae Smith, Kim Labuschagne, Tracy Masebe, Ntanganedzeni Mapholi

**Affiliations:** Department of Agriculture and Animal Health, College of Agriculture and Environmental Sciences, University of South Africa, Roodepoort, South Africa; Department of Life and Consumer Sciences, College of Agriculture and Environmental Sciences, University of South Africa, Roodepoort, South Africa; South African National Biodiversity Institute (SANBI), Foundational Biodiversity Sciences, Pretoria, South Africa

## Abstract

The lion (*Panthera leo melanochaita*) is one of the most iconic species and part of the big five, maintaining ecological balance and a major wildlife-based tourism attraction in South Africa. Despite its importance, it is currently threatened by rapid population decline and increasing population fragmentation. Therefore, there is a need for a high-quality genomic resource that captures the diverse genetic landscape of South African lion populations. To address this, we present a high-quality genome assembly of the lion, generated using PacBio HiFi and Omni-C sequencing technologies. The final assembly comprises 2.45 Gb, with a scaffold N50 of 148 Mb and a contig N50 of 22 Mb. Remarkably, 94.8% of the genome is anchored to 19 scaffolds, reflecting the high degree of contiguity and near-complete chromosomal level. Completeness assessment of the genome showed 98.2% BUSCO, 98.2% k-mer completeness and QV of 65.6 underscoring high accuracy and biological integrity. Genome annotation predicted 831.4 Mb (33.9%) of repetitive sequences and 21 739 protein-coding genes. This work provides high-quality genomic resource to establish a foundation for future population genomic and conservation-focused investigations of the lion populations in South Africa.

## Background and Summary

The lion (*Panthera leo melanochaita*) is one of the most popular hyper-carnivorous species mainly found in the savanna ecosystems of sub-Saharan Africa ^1^. It is a prominent member of South African wildlife species, and is widely recognised for its distinctive morphological attributes and functional traits including considerable agility, muscular strength and the ability to reach high running speeds of up to 80km/h^2 3^. Collectively, these attributes render the lion as a highly effective apex predator in their natural ecosystem. Lions primarily inhabit the savanna grassland ecosystem as dominant predators, and their density is directly proportional to that of prey ^4^. Therefore, they play a critical role in regulating the population of prey and maintaining ecological balance ^5^. Beyond their ecological importance, the lion industry is one of the most lucrative sectors in South Africa, predicted to generate over US$180 million annual through trade and tourism ^6^.

Despite its ecological and economic significance, lions are classified as vulnerable species globally as well as in South Africa under Section 56(1) of National Environmental Management: Biodiversity Act, 2004 (Act No 10 of 2004). This status reflects a rapid decline in the global population size over the past century, leading to its inclusion on the International Union for Conservation of Nature (IUCN) Red List ^7^. While the lion status in South Africa is considered stable, the population in other African regions (central and West Africa) face a high risk of decline, with projection exceeding 50% in the next 20 years ^8^. In the last century, lion populations were estimated at around 200 000 globally, however, the current estimates range between 23 000 to 39 000 free roaming lions, primarily found in Africa and some parts of India ^4,9^.

According to the South African Department of Forestry, Fisheries and the Environment (DFFE) report of 2018, South Africa is estimated to have about 11 155 lions. Of these, about 8000 (72%) are distributed across privately owned and government registered breeding facilities, many of which are driven by commercial and financial interest rather than conservation objectives. The remaining 3 155 (28%) lions roam freely in the Kruger National Park across Mpumalanga and Limpopo provinces, followed by the Kgalagadi Transfrontier Park in the Northern Cape, with smaller numbers found in other national parks ^10^. The decline of lion populations is driven by human and environmental factors, including habitat loss and fragmentation due to human population growth. Legal and illegal hunting activities also play a role in the declining population due to the controversial practice of trophy hunting and poaching motivated by the global demand for lion’s body parts to be used for medicinal and ornamental purposes ^11,12^.

The establishment of privately owned breeding facilities and associated commercial trade has further contributed to the fragmentation of the lion population in South Africa. This fragmentation increases the risk of inbreeding and reduces gene flow among sub-populations ^13-15^. Disruption in gene flow promotes genetic divergence between isolated groups which can erode the population’s genetic structure and challenge its long-term viability ^16^. Given the magnitude of the threats facing South African lions, co-ordinated conservation efforts are urgently needed. The first step is to develop a high-resolution genomic resource that accurately represent the species in the region. Generating genomic resources of a lion enables critical downstream research including comparative and functional genomics. A high-quality and annotated genome will provide valuable insights into the effective population size, population structure and genetic diversity, thereby facilitating precise and effective conservation strategies for the South African lion. Therefore, the current study utilised PacBio HiFi long reads and Omni-C sequencing technologies to generate a high-quality chromosome-level genome assembly of the South African lion.

## Materials and methods

All research procedures were approved by the University of South Africa ethics committee (AREC-100818-024) and conducted in accordance with the relevant guidelines and regulations.

### Genome sample collection

Three EDTA blood samples collected from captive lions were provided by the South African National Biodiversity Institute (SANBI) Wildlife Biobank, Pretoria, South Africa. The collected blood samples were transported for analysis at the laboratory using a cold-chain and were stored in -80 °C freezer before being processed.

### Extraction, library preparation and sequencing

A high molecular weight (HMW) DNA was extracted using 500 μL of blood using Monarch® HMW DNA Extraction Kit for Cells & Blood, following the manufactures’ protocol. The concentration and purity of the DNA was then quantified using Qubit™ dsDNA Quantification Broad Range Assay (Invitrogen, Q32853). To generate high-fidelity (PacBio HiFi) long read data, the library was prepared using SMRTbell® prep kit version 3.026 (Pacific Biosciences). The final library was sequenced using the PacBio Revio platform producing 77.6 Gb data with the average read length of 11.5 kb corresponding to 33.75X coverage. For Omni-C data, the library was prepared using Dovetail Omni-C Kit (Dovetail Genomics) following the manufacturers protocol and then sequenced using PacBio Onso sequencing platform yielding 57.2 Gb of raw data corresponding to 20X coverage.

### Genome assembly

The assembly of the South African lion genome was assembled using the following pipeline: HiFi raw reads were first assessed for quality control using Fastlong v0.3.0 ^17^ to remove adapter sequences and to filter out low-quality reads resulting in 7.2 million reads in with average of 11.5 kb of sequence data. Subsequently, the Omni-C reads also underwent quality assessment using Fastp v0.24.0 ^18^, revealing pair-end reads with high quality. To estimate the genome size, GenomeScope v2.1.0 ^19^ software was used using k-mer profiling with k-mer size of k=21 and estimated a genome size of 2.11 Gb, with homozygosity and heterozygosity levels of 98.9% and 1.08%, respectively, an error rate of 0.0876%.

The assembly was then assembled using Hifiasm v0.20.0 ^20^ software in Hi-C mode. Thereafter primary assembly was then further analysed to the chromosomal level in which Omni-C data was integrated to the following the Dovetail workflow (https://dovetail-analysis.readthedocs.io/en/latest/index.html). In these steps, BWA v0.7.17 tool ^21^ was used to map the Omni-C data to the draft reference genome which removed all low-quality mapped reads. Pairtools v1.0.3 ^22^ was then applied to remove PCR duplicates. The processed alignments used to generate a Hi-C contact matrix with Juicer Tools v1.6 ^23^ was further scaffolded using Yet Another Hi-C Scaffolding (YaHS) tool ^24^ to generate chromosome-level assembly. The final assembly resulted in a genome size of 2.45 Gb with scaffold and contig N50 of 148Mb and 22Mb, respectively (Table 1). In this table, the assembled genome is presented alongside previously published lion assemblies^25,26^. The assembly resulted with 94.8% of the scaffolds corresponding to 19 chromosomal blocks also highly contiguous, representing the full diploid set of Lion Chromosomes (Figure 1A).

**Table 1.** Genome assembly statistics of the South African lion (*Panthera leo melanochaita*) in comparison with other lion assemblies.

| Assembly metrics | South African lion<br>(*Panthera leo melanochaita*) | Reference genome-Liger<br>(GCF_018350215.1)<sup>25</sup> | "Brooke"<br>(GCA_046255005.1)<sup>26</sup> | Kenyan lion<br>(GCA_012313985.1)<br>(Unpublished) |
| --- | --- | --- | --- | --- |
| Genome size (Gb) | 2.45 | 2.30 | 2.40 | 2.40 |
| Number scaffolds | 393 | 53 | 8 060 | 240 216 |
| Scaffold N50 (Mb) | 148 | 147.4 | 136,1 | 0.023 |
| Contig N50 (Mb) | 22 | 77.8 | - | 0.023 |
| Contig L50 | 7 | 7 | 2 481 | - |
| Gaps (Mb) | 0.057 | 0.009 | 18,2 | 0 |
| GC content (%) | 41.8 | 41.5 | 41.5 | 41.5 |
| Coverage X | 34 | 76 | 46 | 82 |
| Assembly level | Chromosome | Chromosome | Chromosome | Contig |
| Sequencing technology | PacBio Revio;<br>Dovetail Omni-C | PacBio Sequel 2 | Illumina; Oxford<br>Nanopore; 10X Genomics | Illumina HiSeq |

**Figure 1.**
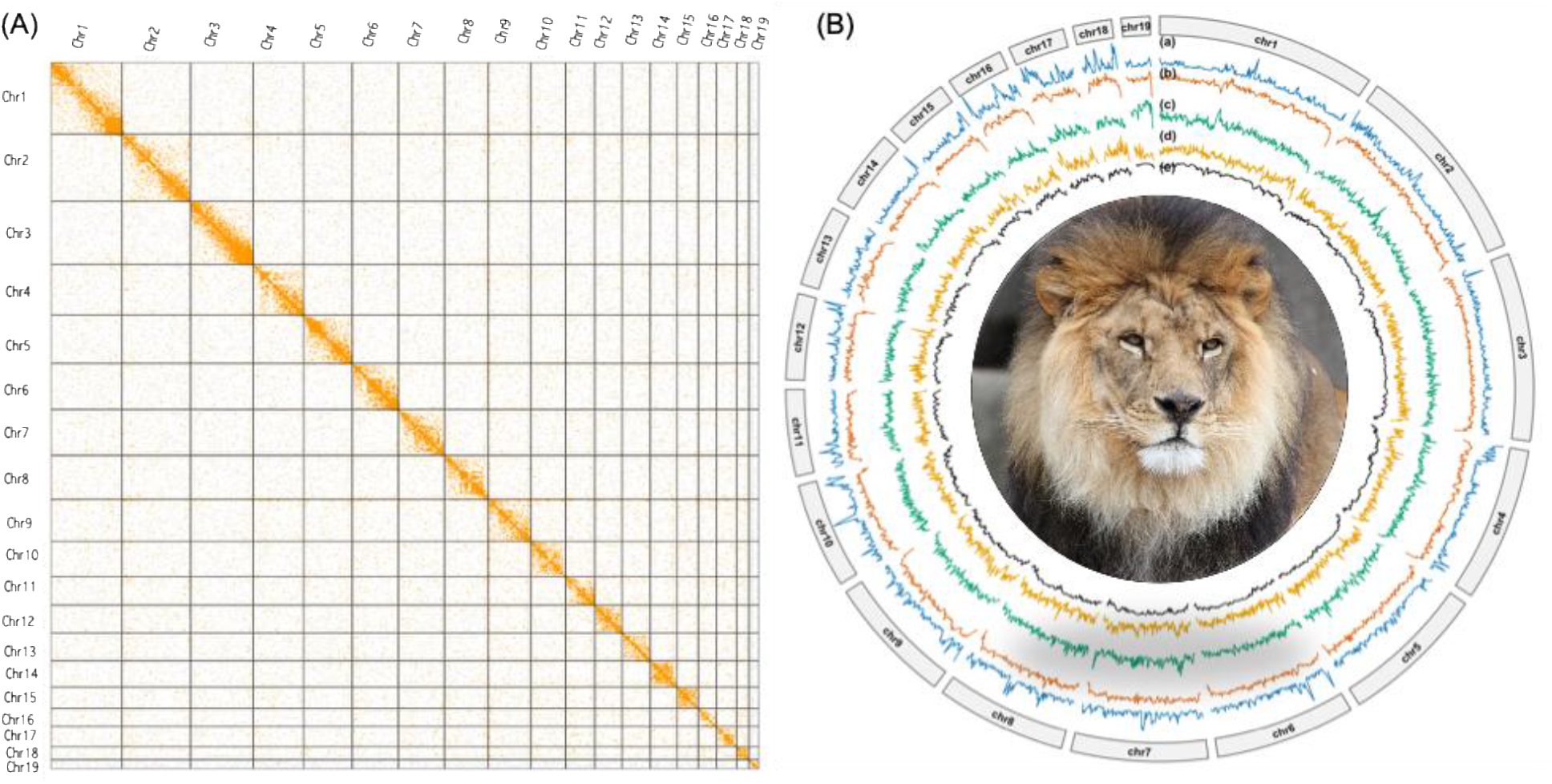
South African genome chromosomal heatmap (A) and Circo’s plot of genome (B) represents genome-wide distributions of major genomic features using 1 Mb non-overlapping windows. From the inner to the outer rings: (a) gene density, (b) total transposable element, (c) long interspersed nuclear element (LINE) density, (d) long terminal repeat (LTR) density and (e) GC content

### Genome assessment

The genome completeness was assessed using Benchmarking Universal Single-Copy Orthologs (BUSCO) using the Carnivora_odb10 database. The assembly captured 98.2% of the conserved orthologs genes present, of which 97% were single copy, 1.2% duplicate, 0.9% fragmented and 0.8% were missing BUSCOs from a total of 14 502 BUSCOs evaluated (Table 2). The current assembly exhibited a higher level of completeness (98.2%) compared with the reference assembly (GCF_018350215.1).

**Table 2.** Genome quality assessment using Benchmarking Universal Single-Copy Orthologs (BUSCO)

| BUSCO categories | Count | BUSCO (%) |
| --- | --- | --- |
| **Complete BUSCOs** | 14 247 | 98.2 |
| **Complete single copy (S)** | 14 069 | 97 |
| **Complete Duplication (D)** | 178 | 1.2 |
| **Fragmented BUSCOs (F)** | 132 | 0.9 |
| **Missing BUSCOs (M)** | 123 | 0.8 |
| **Total BUSCOs searched** | 14 502 | 100 |

### Genome annotation and functional description

Prior to gene prediction, the repeat element was identified and masked in two steps. Firstly, the repeat elements were detected using Repeat modeler v.2.0.5 ^27^ which utilises various algorithms to generate a high-quality library of transposable elements (TE’). Then, the RepeatMasker tool v.4.1.5 ^28^ utilised the library to search for DNA interspersed repeats for soft masking. Repeat element annotation revealed that repetitive elements accounts for 33.88% (831.38Mb) of the assembled lion genome. The most common repetitive element was Long Interspersed Nuclear Elements (LINEs) at 15.9%, followed by Short Interspersed Nuclear Elements (SINEs) at 10.0%, then Long Terminal Repeat retrotransposons (LRT) elements at 3.6%, and the remaining repeat elements are summarised in Table 3 and Figure 2.

**Table 3.** Descriptive statistics of the repeat elements in the genome of the SA lion.

| Repeat element | Number of elements | Total length occupied (Mb) | Percentage (%) |
| --- | --- | --- | --- |
| **LINEs** | 901 400 | 389.0 | 15.9 |
| **SINEs** | 1 271 399 | 245.3 | 10.0 |
| **LRT elements** | 366 281 | 87.9 | 3.6 |
| **DNA elements** | 360 225 | 45.9 | 1.9 |
| **Small RNA** | 1 083 873 | 221.3 | 9.0 |
| **Simple repeats** | 890 392 | 41.4 | 1.7 |
| **Low complexity** | 108 400 | 6.1 | 0.3 |
| **Unclassified** | 20 489 | 15.3 | 0.6 |

**Figure 2.**
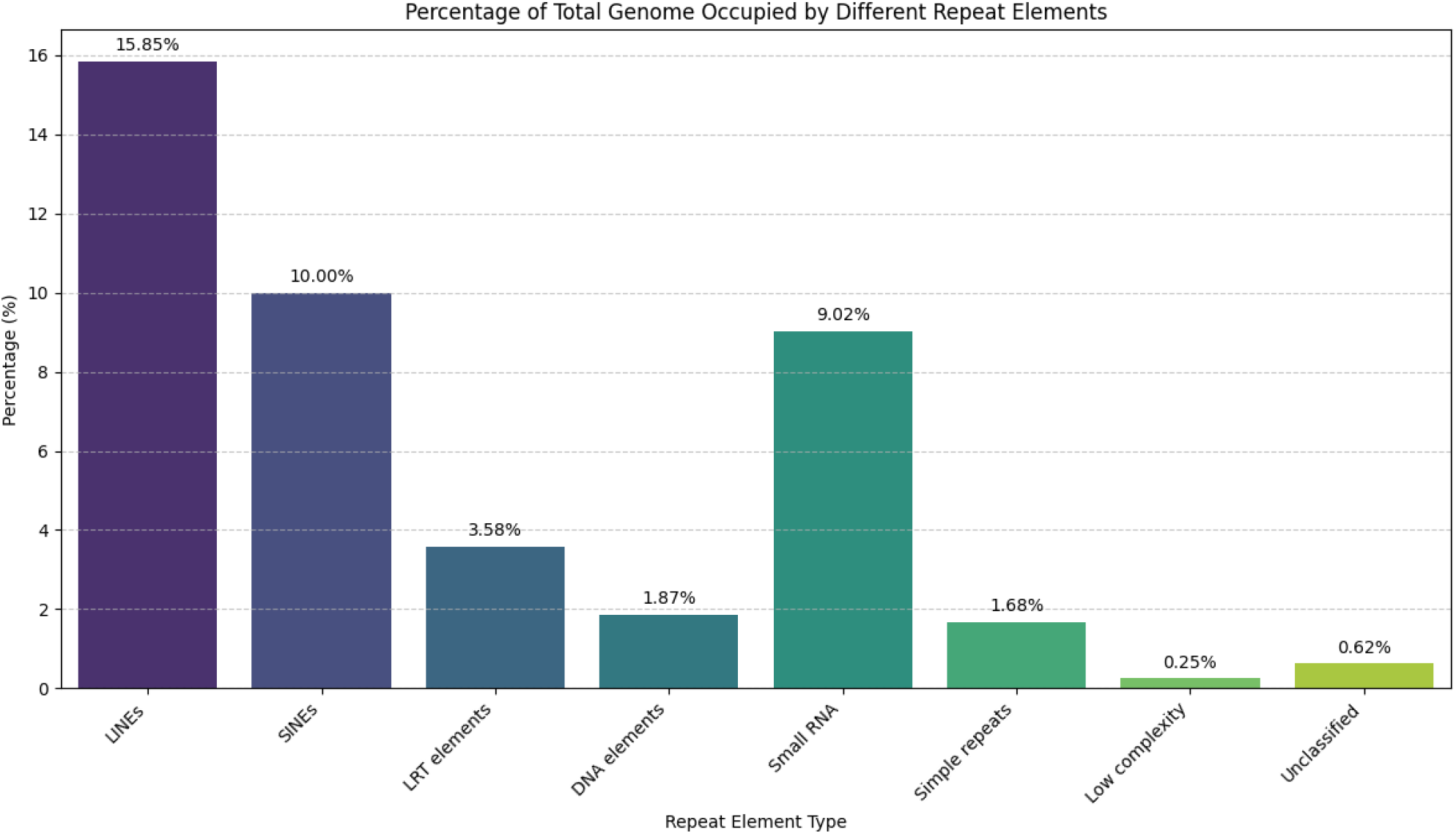
Repetitive elements distribution of the South African lion genome.

Gene prediction was performed on the masked genome using *ab initio* gene prediction tool Tiberius v1.1.4 ^29^, a deep learning gene prediction tool that incorporates external evidence to improve accuracy of the predicted protein-coding genes (Table 4). The annotation identified 21 739 protein-coding genes with a minimum length of 201bp, a maximum length of 74 550 and an average length of 1 450bp. Thereafter, protein genes were assessed for completeness using BUSCO revealed 84.3% completeness.

**Table 4.** Gene structural annotation of the SA lion draft.

| Genome annotation |  |
| --- | --- |
| Protein-coding genes | 21 739 |
| Average gene length (bp) | 1 450 |
| Minimum gene length (bp) | 201 |
| Maximum gene length | 74 550 |
| BUSCO | C:84.3% [S:83.2%, D:1.1%], F:3.9%, M:11.7%, n:14502) |

C-complete, S-complete single-copy, D-complete duplication, F-fragmented. M-missing, and n-total BUSCOs searched.

Functional annotations were carried out using protein genes predicted through eggNOG-mapper v2.1.12 ^30^, which assigns functional information based on ortholog information from the eggNOG database (evolutionary genealogy of genes: Non-supervised Orthologous Groups). Functional annotation assigned 20 975 genes (96.5%) using several protein databases such as Gene Ontology (GO), Kyoto Encyclopaedia of Genes and Genomes (KEGG), Pfam, CAZy, BRITE, and BiGG. These were classified into 147 Clusters of Orthologs Gene (COG) categories. Functional annotation reveals that majority of the analysed genes (n = 6 438; 31%) do not have known or clear biological functions. This highlighted the presence of insufficiently studied genes. The second most prevailing functional category included signal transduction genes (n = 3 871; 18.5%) such as receptors, kinases and other components important for cellular communication and regulatory networks. The third most common category was transcription (COG category K) with 1 834 genes (8.7%) associated with regulating gene expression. The detailed functional annotation results is in the Supplementary table and in Figure 3.

**Figure 3.**
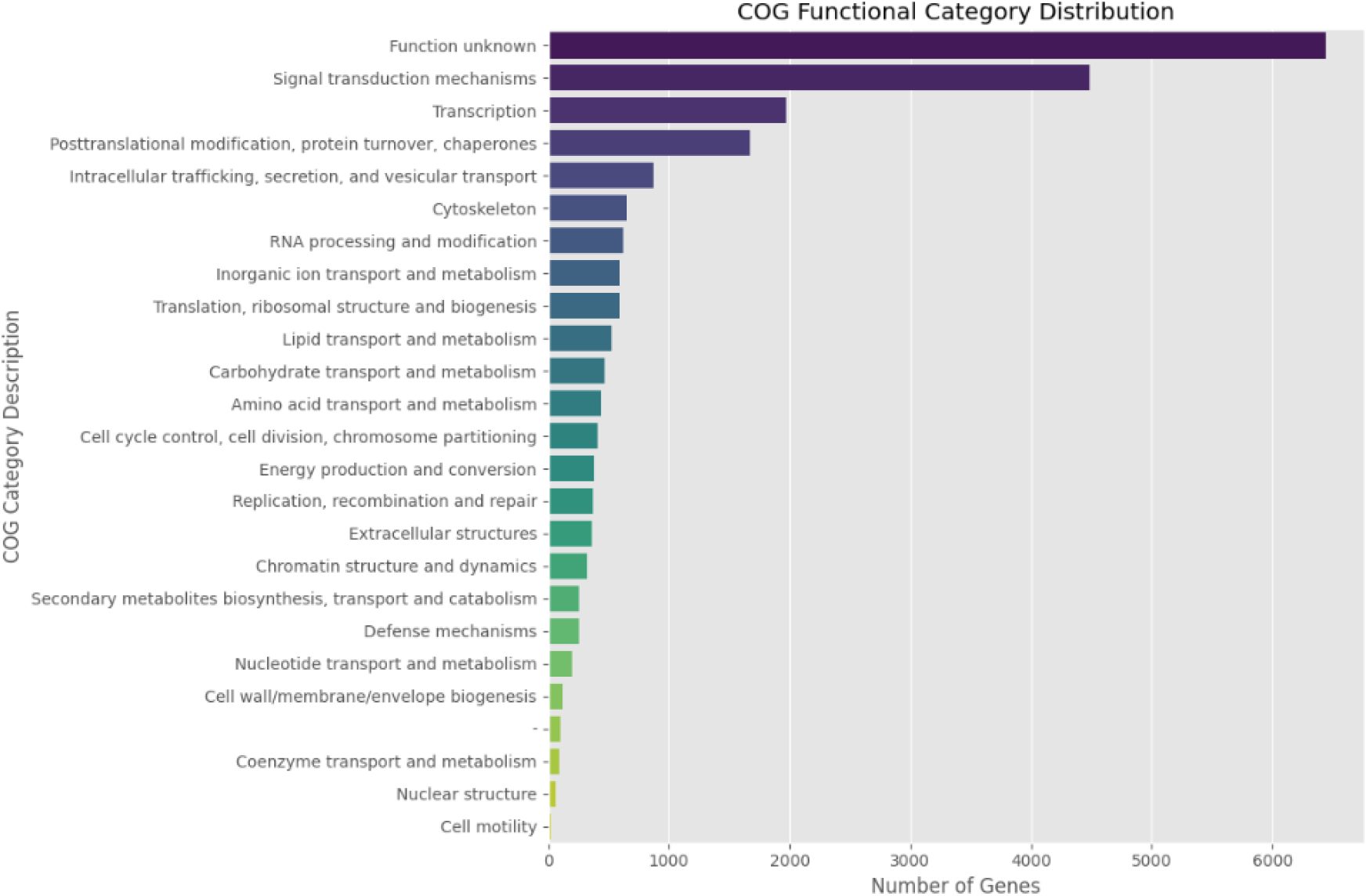
Cluster of Orthologs Gene categories (COG) of the lion.

Gene Ontology (GO) analysis assigned a total of 23 461 GO terms to the lion gene set (Figure 4). The distribution of these annotations across the three principal GO categories revealed that the majority were associated with Biological Process (n = 16 925; 72.1%), followed by Molecular Function (n = 4 486; 19.1%), whereas the smallest proportion was attributed to Cellular Component (n = 2 050; 8.7%).

**Figure 4.**
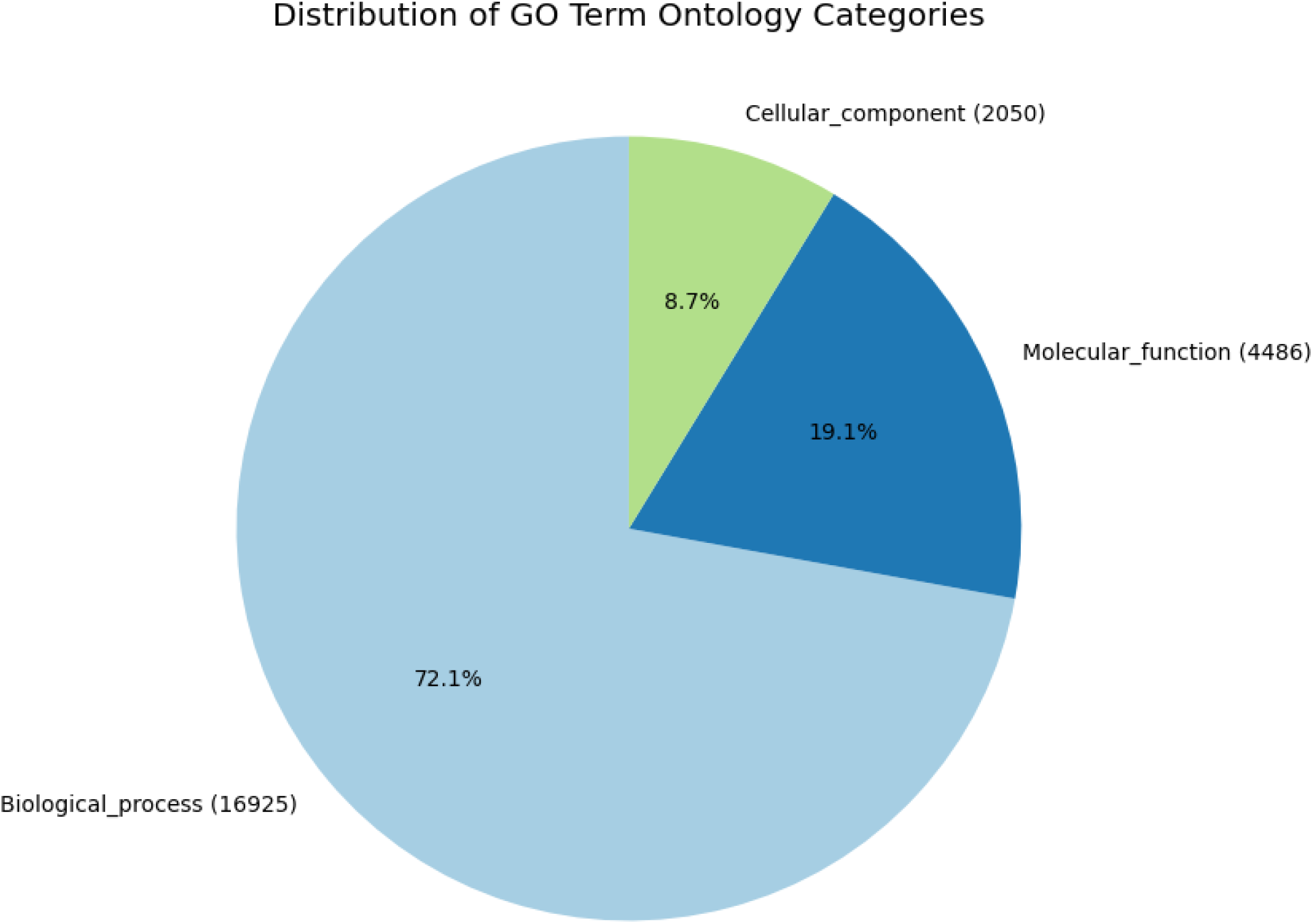
Distribution of Gene Ontology (GO) terms across three principal categories (Biological process, Molecular function and cellular component).

Orthologous assessment among members of the family Felidae was performed on protein fasta files using OrthoFinder v2.5.5. Analysis included protein fasta file of the present lion assembly and for the reference genomes of tiger (GCA_018350195.2), jaguar (GCA_046562885.2), and domestic cat (GCA_018350175.1). Orthologue count per species and the shared orthologues were analysed then visualised using a bar graph and an Upset plot, respectively (Figure 5A-B). The largest number of orthologues (64 526) were observed in domestic cat while the least on the assembled lion (19 546). The analysis identified a set of 15 784 orthologous genes conserved across all four species. A total of 5,850 orthologues were shared exclusively between jaguar, domestic cat, and tiger, then 2 709 orthologues were shared specifically between jaguar and domestic cat. The remaining orthologous relationships are visualised in the Upset plot.

**Figure 5.**
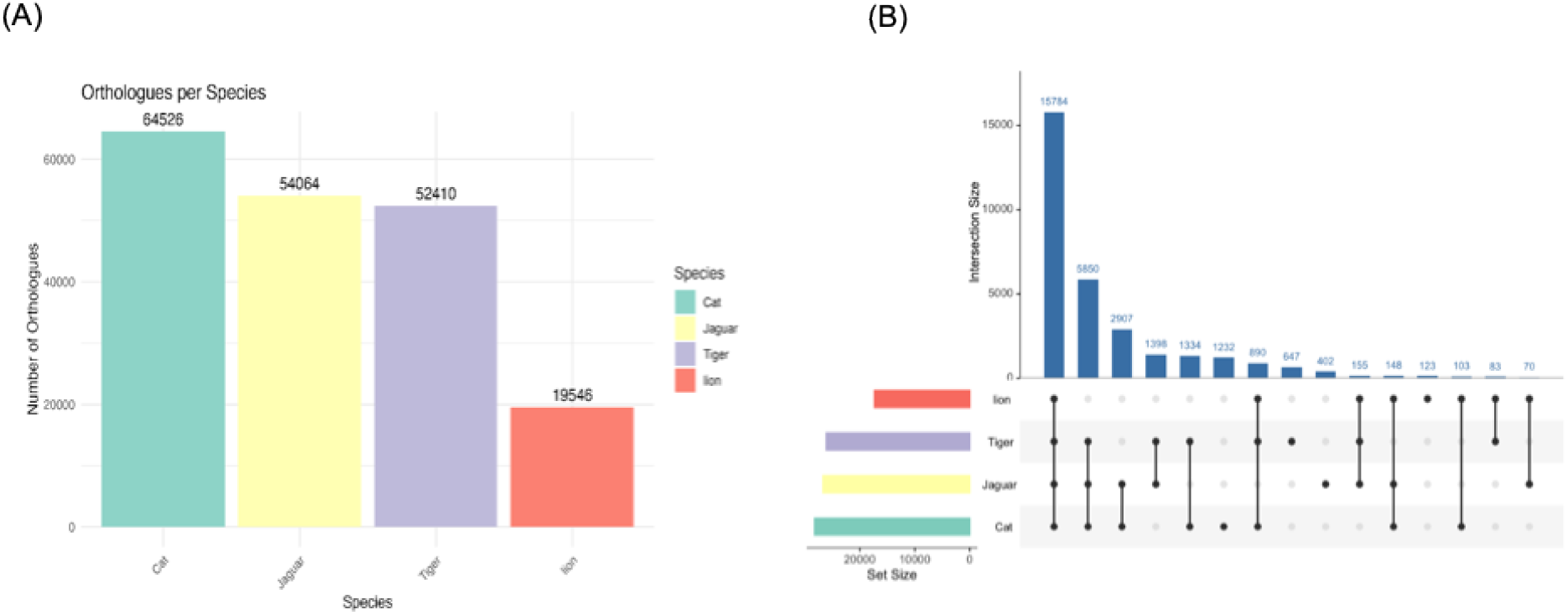
**A-**Distribution of orthologues among Felidae members lion (current assembly), Tiger (GCA_018350195.2), Jaguar (GCA_046562885.2), cat (GCA_018350175.1); **B-**Upset plot showing shared orthologues between members of the Felidae family.

### Data records

All sequencing data generated in this study, including both PacBio HiFi and Hi-C raw reads were deposited to the NCBI database under Bio project accession number: PRJNA1227266, with sequence read archive accessions SRR37525089 and SRR37525088.

### Technical validation

After DNA purification and prior library preparation for both PacBio HiFi and Omni sequencing, DNA underwent quality checks to determine concentration and assessment of integrity. This was done using Qubit™ dsDNA Quantification Broad Range Assay (Invitrogen, Q32853) and a spectrophotometer with HS Assay Kits following the manufacturer’s guide. In this protocol, a 200µl working solution (1 µL of fluorescent dye + 199 µL of buffer) was prepared. Then, 1 µL of DNA was mixed with 199 µL of the working solution and incubated for 2 minutes at a room temperature. Qubit fluorometer measured the florescence and estimated DNA concentration. To evaluate the integrity of the DNA, Agilent Femto Pulse system and the Genomic DNA 50 kb kit (HS 50 kb) were utilised following the manufacturers’ protocol. This process generated electropherogram and the DNA fragment size distribution.

High molecular weight (HMW) genomic DNA from the sample was mechanically sheared to a target fragment size of 15–25 kb using a Megaruptor 3 instrument to generate fragments suitable for PacBioHiFi sequencing. The sheared DNA was subsequently purified using SMRTbell beads, followed by DNA damage repair and end-preparation, and then adapter ligation according to the manufacturer’s protocol. The resulting library was subjected to size selection with a BluePippin system (Sage Science) to remove fragments shorter than 10 kb. The fragment size distribution of the final library was evaluated using a Fragment Analyzer, after which the library was sequenced on a PacBio Sequel II platform. To generate Omni-C data, the same sample used for PacBioHiFi sequencing was utilised to in the preparation of the Dovetail Omni-C proximity ligation library following the manufactures protocol.

## Data availability

The genome assembly and its annotation files were deposited to the NCBI database under submission number ID SUB16048291. The data will be accessible through the genome database up on completion of NCBI under the following reference: JBWAXN000000000. For reviewers, the assembled genome and its annotation files are shared on fig share under the following link: 10.6084/m9.figshare.31573471.

## Code availability

All bioinformatic tools and pipelines utilised in the lion assembly were implemented based on the order they are presented in the material and method section, and adopted from the in-house methodology, and no custom codes were utilised.

## Author contributions

Conceptualization: SH,TST, LTN, NM; Data Curation: SH, SM, TST, AM; Formal Analysis: SH, TST; Funding Acquisition: NM, TM; Investigation: SH, TST, LTN; Methodology: SH, TST, NH, RS; Project Administration: NM, TM; Resources: LTN, SH, Software: SH, TST; Validation: SH, LTN, TST, RS; Visualisation: SH, TST; Writing of the original draft: SH; Reviewing and editing of the manuscript: SH, TST, SM, LTN, NH, RS, AM, NM, TM, KL.

## Acknowledgements

We would like to express our gratitude to the University of South Africa for the provision of financial support and the collaboration with African BioGenome project. Special thanks to the Inqaba Biotech laboratory for collaboration on sequencing platform.

## References

1 Riggio, J. et al. The size of savannah Africa: a lion’s (Panthera leo) view. Biodiversity and Conservation 22, 17–35 (2013).

2 Thurman, M. L. How Do Lions Hunt?

3 Vaughan, D. The fastest animals on Earth.

4 Trinkel, M. & Angelici, F. M. in Problematic wildlife: A cross-disciplinary approach 45–68 (Springer, 2015).

5 Everatt, K. T., Moore, J. F. & Kerley, G. I. Africa’s apex predator, the lion, is limited by interference and exploitative competition with humans. Global Ecology and Conservation 20, e00758 (2019).

6 Harvey, R. G. Towards a cost-benefit analysis of South Africa’s captive predator breeding industry. Global Ecology and Conservation 23, e01157 (2020).

7 Bauer, H., Packer, C., Funston, P., Henschel, P. & Nowell, K. (2016).

8 Bauer, H. et al. Lion (Panthera leo) populations are declining rapidly across Africa, except in intensively managed areas. Proceedings of the National Academy of Sciences 112, 14894–14899 (2015).

9 Riggio, J. et al. Lion populations may be declining in Africa but not as Bauer et al. suggest. Proceedings of the National Academy of Sciences 113, E107–E108 (2016).

10 Department of Environmental Affairs, S. A. Non-detriment finding assessment for Panthera leo (African lion). (2018).

11 Williams, S. T., Williams, K. S., Lewis, B. P. & Hill, R. A. Population dynamics and threats to an apex predator outside protected areas: implications for carnivore management. Royal Society Open Science 4, 161090 (2017).

12 Coals, P. et al. Commercially-driven lion part removal: What is the evidence from mortality records? Global Ecology and Conservation 24, e01327 (2020).

13 Tensen, L., Groom, R. J., Khuzwayo, J. & Jansen van Vuuren, B. The genetic tale of a recovering lion population (Panthera leo) in the Savé Valley region (Zimbabwe): A better understanding of the history and managing the future. PLoS One 13, e0190369 (2018).

14 Trinkel, M. et al. Translocating lions into an inbred lion population in the Hluhluwe-iMfolozi Park, South Africa. Animal Conservation 11, 138–143 (2008).

15 Trinkel, M. et al. Inbreeding and density-dependent population growth in a small, isolated lion population. Animal Conservation 13, 374–382 (2010).

16 Bruche, S. et al. A genetically distinct lion (Panthera leo) population from Ethiopia. European Journal of Wildlife Research 59, 215–225 (2013).

17 Chen, S. Ultrafast one-pass FASTQ data preprocessing, quality control, and deduplication using fastp. Imeta 2, e107 (2023).

18 Chen, S., Zhou, Y., Chen, Y. & Gu, J. fastp: an ultra-fast all-in-one FASTQ preprocessor. Bioinformatics 34, i884–i890 (2018).

19 Ranallo-Benavidez, T., Jaron, K. & Schatz, M. (2020).

20 Cheng, H., Concepcion, G. T., Feng, X., Zhang, H. & Li, H. Haplotype-resolved de novo assembly using phased assembly graphs with hifiasm. Nature methods 18, 170–175 (2021).

21 Li, H. Aligning sequence reads, clone sequences and assembly contigs with BWA-MEM. arXiv preprint arXiv:1303.3997 (2013).

22 Open2C et al. Pairtools: from sequencing data to chromosome contacts. PLOS Computational Biology 20, e1012164 (2024).

23 Durand, N. C. et al. Juicer provides a one-click system for analyzing loop-resolution Hi-C experiments. Cell systems 3, 95–98 (2016).

24 Zhou, C., McCarthy, S. A. & Durbin, R. YaHS: yet another Hi-C scaffolding tool. Bioinformatics 39, btac808 (2023).

25 Chen, W. et al. Whole genome sequencing informs SNP-based breeding strategies to safeguard genetic diversity in captive African lions. Frontiers in Veterinary Science 12, 1577726 (2025).

26 Barazandeh, M., Kriti, D., Fickel, J. & Nislow, C. The Addis Ababa Lions: whole-genome sequencing of a rare and precious population. Genome Biology and Evolution 16, evae021 (2024).

27 Flynn, J. M. et al. RepeatModeler2 for automated genomic discovery of transposable element families. Proceedings of the National Academy of Sciences 117, 9451–9457 (2020).

28 Smit, A., Hubley, R. & Green, P. RepeatMasker Open-4.0 http://www.repeatmasker.org.RMDownload.html 251, p252 (2013).

29 Gabriel, L., Becker, F., Hoff, K. J. & Stanke, M. Tiberius: end-to-end deep learning with an HMM for gene prediction. Bioinformatics 40, btae685 (2024).

30 Cantalapiedra, C. P., Hernández-Plaza, A., Letunic, I., Bork, P. & Huerta-Cepas, J. eggNOG-mapper v2: functional annotation, orthology assignments, and domain prediction at the metagenomic scale. Molecular biology and evolution 38, 5825–5829 (2021).

